# Circular RNA hsa_circ_0068033 inhibits proliferation and invasion of breast cancer cells through sponging miR-659

**DOI:** 10.1101/725523

**Authors:** Pengfei Yuan, Liangliang Lei, Dechun Liu

**Affiliations:** Department of Gastrointestinal Surgery (Breast Surgery), The First Affiliated Hospital, and College of Clinical Medicine of Henan University of Science and Technology, Luoyang, 471003, China

**Keywords:** circular RNA, hsa_circ_0068033, breast cancer, miR-659

## Abstract

Recently, dysregulated circular RNAs (circRNAs) have been associated with the progression of numerous malignant tumors. However, the mechanism through which circRNAs participate in breast cancer (BC) remains unclear. This study was designed to illustrate the role of hsa_circ_0068033 in BC. A series of functional experiments were conducted to assess the function of hsa_circ_0068033 in the development of BC and the underlying mechanisms. The results suggested that the expression of hsa_circ_0068033 was downregulated in BC tissues, and markedly correlated with tumor size (*P*=0.021) and the Tumor, Node, and Metastasis stage (*P*=0.023). Receiver operating characteristic analysis showed that hsa_circ_0068033 testing yielded an area under the curve value of 0.8480 in discriminating BC from non-tumor controls. Functionally, overexpression of hsa_circ_0068033 could inhibit the growth, clone formation, invasion, and migration of MCF-7 and MDA-MB-231 cells, while activating the intrinsic apoptotic pathway to induce apoptosis. The xenograft experiment revealed that exogenous hsa_circ_0068033 is able to reduce the tumorigenic ability of MDA-MB-231 cells in nude mice. Rescue assays further proved that hsa_circ_0068033 exerts biological functions by sponging miR-659. This study revealed for the first time that hsa_circ_0068033 acts as a tumor suppressor gene in BC, and the hsa_circ_0068033/miR-659 axis participates in the progression of BC.

## 1. Introduction

Breast cancer (BC) is one of the most common malignant tumors in females worldwide, and its incidence is increasing on an annual basis [1]. According to the latest cancer statistics, BC has the highest incidence and the second highest mortality among all tumors in female patients [2]. The occurrence, development, invasion, and metastasis of BC involve a complicated process in which multiple genes work in synergy. This process is associated with the abnormal expression of many genes, including the activation of oncogenic genes, deactivation of tumor suppressor genes, etc. [3]. Therefore, discovering the mechanism of BC development from the perspective of gene abnormality may help understand the evolutionary process of BC, and have great implications for its diagnosis, treatment, and prediction of prognosis.

Circular RNA (circRNA), a class of endogenous non-coding RNAs different from linear RNA molecules, does not include a 5’ end cap structure and 3’ end poly (A) tail [4, 5]. It is a closed circular structure formed by covalent bond and widely present in eukaryocytes [5]. CircRNA was initially thought to be a splicing intermediate, byproduct, or an accidental splicing error after transcription. Along with the rapid development of high-throughput sequencing and bioinformatics, an increasing number of cirRNAs have been found in organisms, and their biological functions have been further understood [5]. Currently, it has been confirmed that cirRNA can regulate the expression of its target gene via a micro RNA (miRNA) sponge, and that it can adjust pre-mRNA splicing to influence protein production [6, 7]. In addition, circRNA can directly bind to proteins, or establish an indirect RNA-mediated association with proteins to influence their functions [8]. Studies have reported that circRNA is closely correlated with the occurrence and development of various malignant tumors, including liver cancer [9, 10], lung cancer [11, 12], cervical cancer [13] gastric cancer [14, 15], colorectal cancer [16, 17], etc. In recent years, the potential applications of circRNA in the regulation of biological functions and the clinical diagnosis and prognosis of BC have received considerable attention. Our previous studies have shown that the expression of hsa_circ_0068033 was decreased in certain patients with BC. It is speculated that hsa_circ_0068033 may act as a tumor suppressor gene. Based on previous evidence, this study was designed to investigate the expression of hsa_circ_0068033 in tissues obtained from patients with BC, and its potential value in the clinical diagnosis of BC. Furthermore, we sought to explore the biological functions of hsa_circ_0068033 in BC both *in vivo* and *in vitro*, and trace the underlying molecular mechanisms.

## 2. Materials and methods

### 2.1. Patient specimens

Tumor tissues (n=36) and paired adjacent normal tissues were collected from BC patients who were treated at the First Affiliated Hospital, and College of Clinical Medicine of Henan University of Science and Technology. All tissue specimens were obtained through surgery or biopsy for the first time and confirmed by histopathological examination. Immediately after collection, the tissue specimens were cryopreserved at −80°C. The collection of all the specimens was approved by the Ethics Committee of the Hospital.

### 2.2. Cell culture and transfection

All BC cell lines were puchased from the Chinese Academy of Sciences (Shanghai, China). MCF10A, MCF-7, T47D, and MDA-MB-468 cells were cultured in Dulbecco’s Modified Eagle’s Medium high-glucose medium or RPMI-1640 medium. Both media contained 10% fetal bovine serum (FBS). MDA-MB-231 cells were cultured in L-15 culture solution containing 10% FBS. The sequence of hsa_circ_0068033 was obtained from the University of California, Santa Cruz database (http://genome.ucsc.edu/index.html), cloned into a PLCDH-cir vector (PLCDH-cirR0068033), and subsequently packaged in a lentivirus. Mimics of miR-659 (including negative controls) were synthesized (GenePharma) and transfected to the target cells using Lipofectamine™ 2000 (Invitrogen). The experiment included three groups, namely the overexpression group (Oe-circRNA), mock-vehicle group (Oe-vector), and Oe-circRNA + mimic group.

### 2.3. Quantitative reverse transcription-polymerase chain reaction (qRT-PCR)

Total RNA was extracted with the RNAprep pure Tissue Kit (TIANGEN, China), and reversely transcribed into cDNA with 0.5–1 μg RNA as the template. Extraction of nuclear RNA was performed using the PARIS™ Kit (Life technologies). The PCR system (20 μL in total) contained 2×SYBR Green Mix (Roche) 10 μL, cDNA 1 μL, upstream and downstream primer 5 pmol; ddH_2_O was added until the total volume reached 20 μL. Three replicates were performed for each specimen, and *U6* and *GAPDH* served as internal references. The 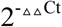 was calculated based on the cycle threshold (Ct) value of product amplification for relative quantification.

### 2.4. In situ hybridization (ISH)

ISH was performed mainly using the Double-DIG labeled circRNA ZKSCAN1 probe Kit (Exiqon). The major steps were as follows: tissue slice de-waxing, de-proteinization, pre-hybridization, hybridization, flushing, sealing, color development with BCIP/NBT, nuclear staining with nuclear fast red solution, dehydration, and mounting.

### 2.5. Cell viability assay

Cells in the logarithmic phase were inoculated into 96-well plates (1×10^3^ cells/well). There were four repeats in each group. The Cell Counting Kit-8 (CCK-8; Dojindo, Japan) reagent was added (1/10 of the total volume) at day 0, 1, 2, 3, and 4 after culture, and the cells were further cultured for 3 h. The optical density value in each well (492 nm/450 nm) was measured using a microplate reader (Thermo).

### 2.6. Colony formation assay

Cells were inoculated into six-well plates at a density of 800 cells/well. These plates were placed in an environment with saturated humidity at 37°C/5% carbon dioxide prior to stilling culture for 2–3 weeks. Culture was continued until cell clones in the culture dish were visible to the naked eye. Acetic acid/methanol (1:3) (5 mL) was introduced for fixation for 15 min. After staining with a proper amount of Giemsa stain for 10–30 min, clones with >50 cells were counted under a low-power microscope (OLYMPUS).

### 2.7. Wound healing migration assay

Approximately 5×10^5^ cells were added to each well of the six-well plate, for the overnight fusion rate to reach >90%. The following day, a vertical scratch was made using a 200-μL pipette tip against a straight ruler. After rinsing thrice with phosphate-buffered saline, the scratched cells were removed. Images were captured to calculate the number of migrated cells at 0, 24, and 48 h.

### 2.8. Transwell assay

Matrigel was diluted with FBS-free basal medium. Briefly, 1×10^5^ cells/well were inoculated into the upper Transwell chamber, and 800 μL culture solution containing 10% FBS was added into the lower Transwell chamber for chemotaxis. Subsequently, the 24-well plate was placed in an incubator for 24 h at 37°C/5% carbon dioxide. The cells and Matrigel in the Transwell chamber were scraped using a filter paper, fixed in 4% paraformaldehyde for 10 min, and stained with 0.1% crystal violet for 30 min. Photos were captured under an optical microscope (OLYMPUS). The number of cells was counted in at least five fields to calculate the mean value.

### 2.9. Terminal deoxynucleotidyl transferase UTP nick end labeling (TUNEL) and DNA ladder assay

The apoptosis experiment was conducted using the Colorimetric TUNEL Apoptosis Assay Kit and Apoptosis (Beyotime, China), and DNA Ladder Extraction Kit (Beyotime, China) with a Spin Column according to the instructions provided by the manufacturer.

### 2.10. Immunoblotting

Total protein was extracted using radioimmunoprecipitation assay buffer (RIPA Lysis Buffer; Beyotime, China), and the protein concentration was determined through the bicinchoninic acid method (Beyotime, China). After electrophoretic separation with 12% separation gel at a constant voltage of 80 V, the proteins were transferred to a polyvinylidene difluoride membrane at a constant current of 250 mA. Subsequently, tris-buffered saline and Tween 20 containing 5% skim milk powder was added to the membrane for blocking overnight. Rabbit anti-human monoclonal primary antibody B-cell lymphoma-2 (Bcl-2) (Abcam, 1:1,000) and mouse anti-human GAPDH monoclonal primary antibody (Abcam, 1:1,000) were added for incubation at room temperature for 4 h. Subsequently, horseradish peroxidase-labeled goat anti-mouse/rabbit (Abcam, 1:10,000) secondary antibody solution was added for 2 h at room temperature. Enhanced chemiluminescence reagent (Beyotime, China) was added for film development and photography.

### 2.11. Xenograft experiment

The concentration of BC cells in each group was 2×10^7^ cells/mL. Female BALB/C nude mice were subcutaneously inoculated with the cells on the back (0.2 mL/mouse). Approximately 10 days later, palpable rice-sized nodules developed at the site of inoculation, and the experiment was initiated at this point. Tumor volume and weight were measured once weekly and 28 days after the experiment, respectively. The experiment was approved by the Ethics Committee of the First Affiliated Hospital, and College of Clinical Medicine of Henan University of Science and Technology.

### 2.12. Dual luciferase reporter assay

MiRNA molecules interacting with hsa_circ_0068033 and binding sites were predicted by the Circular RNA Interactome (https://circinteractome.nia.nih.gov/). The sequence of wild-type hsa_circ_0068033 containing potential miR-659-binding sites was cloned into a pmirGLO vector downstream of luciferase separately (cirRNA-wt group). In addition, a mutant hsa_circ_0068033 sequence was designed to connect with the pmirGLO vector (cirRNA-mut group). The recombinant vector and miR-659 mimic were jointly transfected into MDA-MB-231 cells using Lipofectamine™2000 (Invitrogen). These cells were lysed after 24 h of culture and the activity of luciferase was determined.

### 2.13. Statistical analysis

SPSS 16.0 software (IBM Corporation, Armonk, NY, USA) was used for stastical analysis. Mesurement data of the experiement results were expressed as Mean±SD. All the data of this study were obtained after repeating experiment at least 3 times. If normality and homogeneity of variance were met, the differences between two groups were compared by Student’s *t*-test. The χ^2^ test or corrected χ^2^ test was used to analyze the association between hsa_circ_0068033 expression and the clinical features. P<0.05 was considered to suggest statistical significance.

## 3. Results

### 3.1. Hsa_circ_0068033 was downregulated in BC tissues

The expression levels of hsa_circ_0068033 in 36 pairs of BC tissues and normal adjacent tissues were detected using qRT-PCR. The results showed that the expression of hsa_circ_006803 in BC tissues was obviously lower than that measured in normal adjacent tissues (*P*<0.001) (Figure 1A and 1B). Moreover, ISH revealed that positive expression of hsa_circ_0068033 could be detected in normal adjacent tissues, whereas it was negative in BC tissues (n=3) (Figure 1C). When grouped according to clinical staging, the results suggested that the expression of hsa_circ_0068033 in stage I–II BC patients was increased versus that reported in stage III–IV BC patients (*P*<0.05) (Figure 1D). Furthermore, clinicopathological association analysis showed that the expression of hsa_circ_0068033 was correlated with tumor size (*P*=0.021) and the Tumor, Node, and Metastasis stage (*P*=0.023) (Table 1), indicating that hsa_circ_0068033 may participate in the progression of BC. Additionally, we examined five cell lines, and the findings showed that the expression levels of hsa_circ_0068033 in four cell lines were down-regulated compared with those reported in MCF10A normal human mammary epithelial cells. Notably, the relative expression was the lowest in MCF-7 and MDA-MB-231 cells (*P*<0.05) (Figure 1E); thus, these cell lines were used for the subsequent experiments. Receiver operating characteristic (ROC) analysis revealed that the area under the curve (AUC) of hsa_circ_0068033 in distinguishing tumor from non-tumor reached 0.8480 (*P*<0.01), suggesting a relatively high diagnostic value (Figure 1F).

**Table 1.** Clinical characteristics of circRNA hsa_circ_0068033 expression in BC

| Clinicopathological features | Low (n=20) | High (n=16) | $\chi^2$ | P value |
| --- | --- | --- | --- | --- |
| Age (years) |  |  |  |  |
| ≤50 | 8 | 7 | 0.051 | 0.821 |
| >50 | 12 | 9 |  |  |
| Tumor size (cm) |  |  |  |  |
| ≤2 | 6 | 11 | 5.335 | 0.021 |
| >2 | 14 | 5 |  |  |
| TNM stage |  |  |  |  |
| I-II | 5 | 10 | 5.143 | 0.023 |
| III-IV | 15 | 6 |  |  |
| Lymphatic metastasis |  |  |  |  |
| Yes | 9 | 7 | 0.006 | 0.94 |
| No | 11 | 9 |  |  |

**Figure 1.**
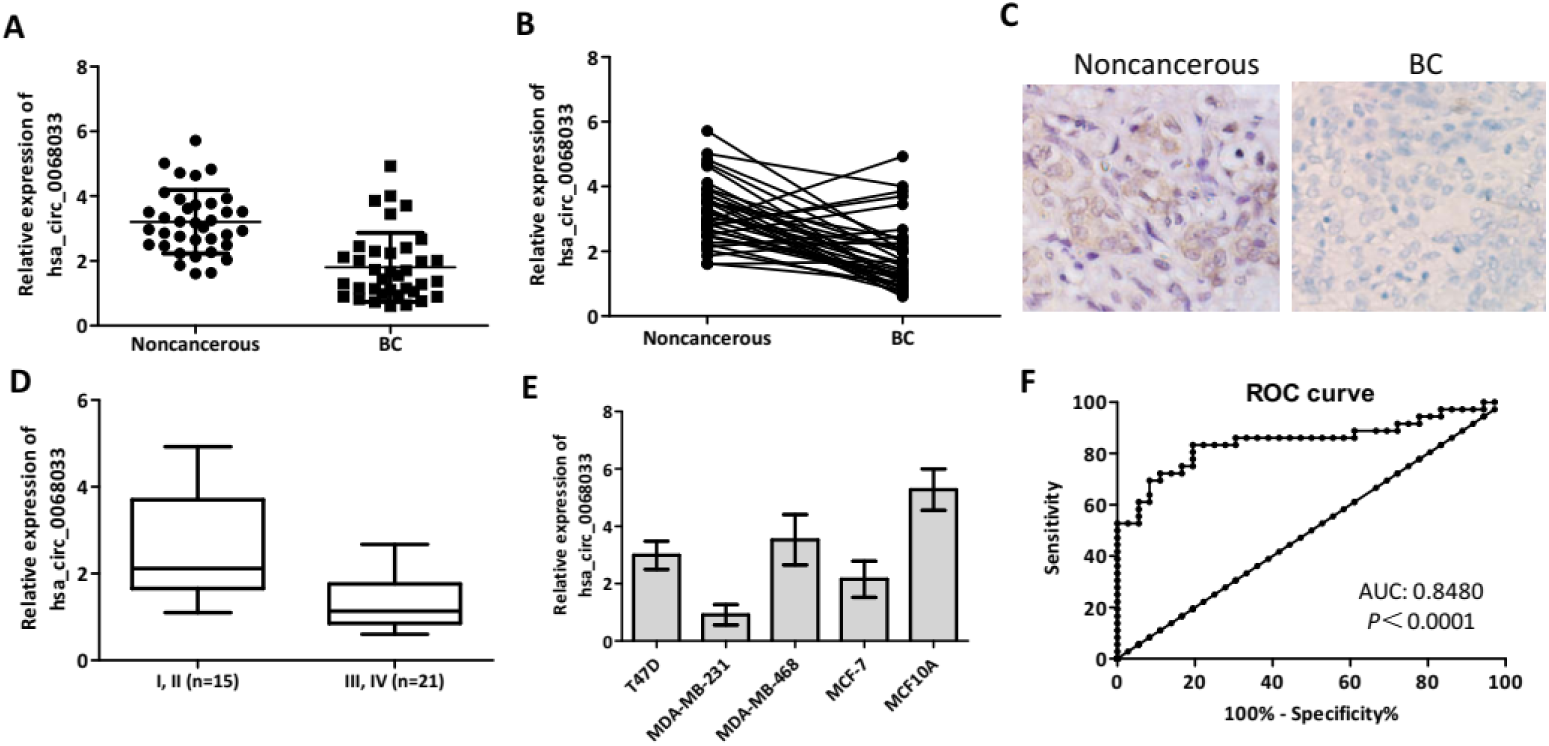
Expression of hsa_circ_0068033 in BC tissues and cell lines. (A, B) The qRT-PCR results showed that hsa_circ_0068033 was downregulated in BC tissues (n=36) compared with adjacent non-cancerous tissues (n=36). (C) ISH showed positive expression of hsa_circ_0068033 in adjacent non-cancerous tissues, and negative expression in BC tissues (n=3). (D) The levels of hsa_circ_0068033 were lower in stage III–IV BC patients versus stage I–II patients. (E) The qRT-PCR analysis showed that hsa_circ_0068033 was downregulated in BC cell lines. (F) The plotted ROC curve of hsa_circ_0068033 in confirming BC (AUC=0.8480, *P*<0.0001).

### 3.2. Overexpression of hsa_circ_0068033 inhibited proliferation and invasion in BC cells

After transfection with a recombinant lentiviral vector (PLCDH-cirR0068033), the expression levels of hsa_circ_0068033 in MCF-7 and MDA-MB-231 cells were significantly increased (Figure 2A). The result of CCK-8 assay suggested that overexpressed hsa_circ_0068033 could inhibit the growth of MCF-7 and MDA-MB-231 cells (Figure 2B). The clone formation experiment showed that the numbers of cloned MCF-7 and MDA-MB-231 cells in the Oe-circRNA group were 79.75±11.38 and 86.50±11.68, respectively. These numbers were markedly lower than those observed in the Oe-vector group (119.00±13.32 and 108.75±6.90 respectively; P<0.05 for all) (Figure 2C). In addition, the numbers of migrated and invasive cells in the Oe-circRNA group were lower than those reported in the Oe-vector group (Figure 2D and 2E), indicating that overexpressed hsa_circ_0068033 may reduce the invasion and migration of MCF-7 and MDA-MB-231 cells. The results of the aforementioned experiment showed that exogenous hsa_circ_0068033 may suppress the growth, proliferation, invasion, and migration of MCF-7 and MDA-MB-231 cells *in vitro*. Furthermore, we further verified these findings *in vivo*. As shown in the experiment of tumor formation in nude mice, the Oe-circRNA group had a much lower tumor volume and weight and a significantly reduced tumorigenic ability than the Oe-vector group in MDA-MB-231 cells (Figure 2F). Intriguingly, there was no tumor formation in the nude mice of the Oe-circRNA and Oe-vector groups after injection with MCF-7 cells (Figure 2F).

**Figure 2.**
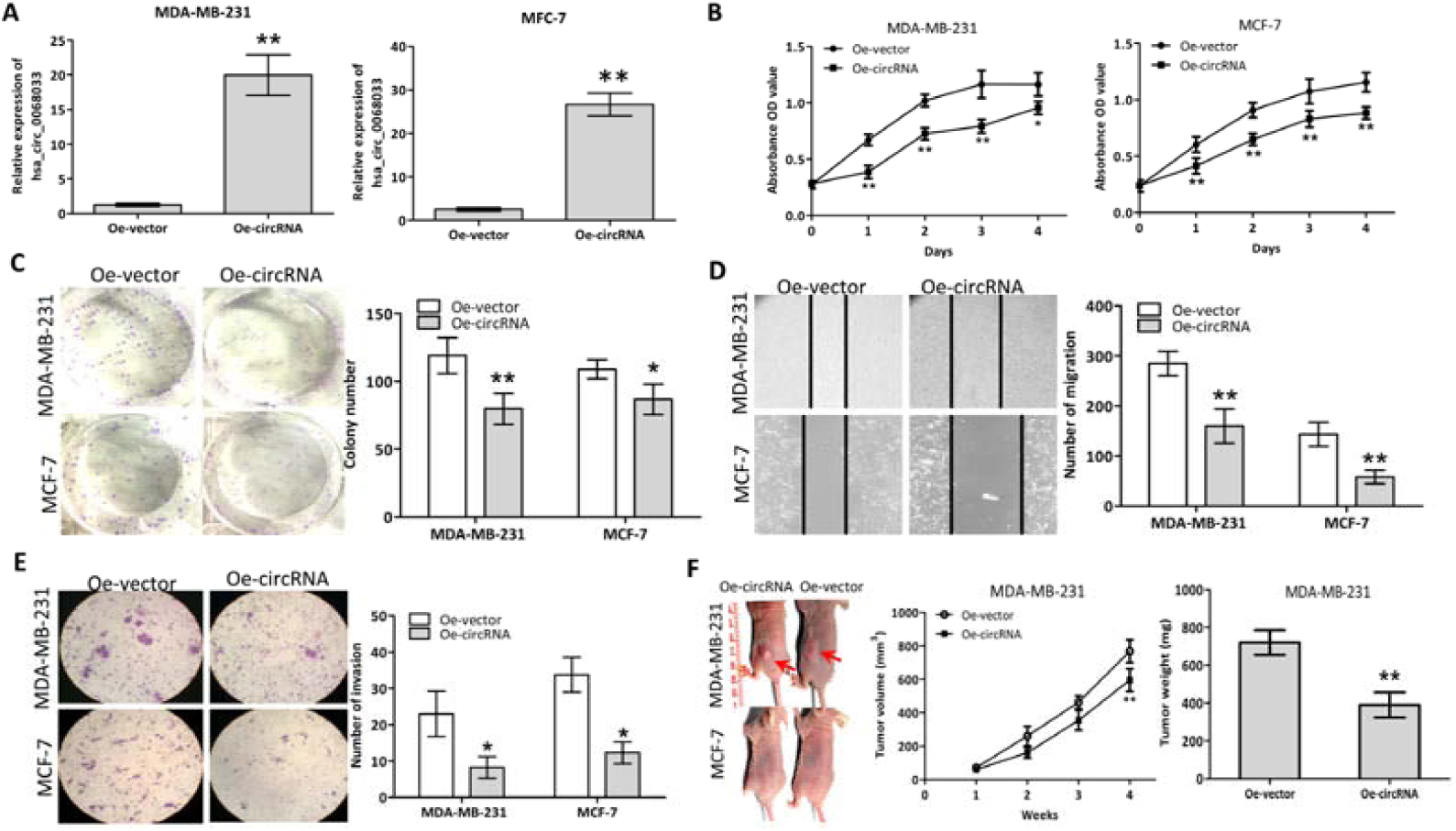
Exogenous hsa_circ_0068033 suppresses the growth, proliferation, invasion, and migration of MCF-7 and MDA-MB-231 cells. (A) The levels of hsa_circ_0068033 were markedly increased following cell transfection with PLCDH-cirR0068033. (B) Cell viability assay using CCK-8. (C) Colony formation assay. (D) Wound healing migration assay. Migrated cells were counted at 48 h after transfection. (E) Transwell invasion assay. Cells were at counted 48 h following transfection with PLCDH-cirR0068033. (F) Xenograft experiment. MDA-MB-231 and MCF-7 cells transfected with PLCDH-cirR0068033 were injected into the back of BALB/C nude mice. Tumor volume was monitored once weekly for 4 weeks. Tumor weights were measured at the endpoint of the xenograft experiment. Red arrows indicate the tumors. \**P*<0.05, \*\**P*<0.01.

### 3.3. Overexpression of hsa_circ_0068033 promoted apoptosis in BC cells

We further validated the potential apoptosis-inducing effect of exogenous hsa_circ_0068033 in MCF-7 and MDA-MB-231 cells. The TUNEL analysis revealed that the numbers of apoptotic MCF-7 and MDA-MB-231 cells in the Oe-circRNA group were significantly higher than those recorded in the Oe-vector group (Figure 3A). Following transfection, an obvious DNA ladder appeared in the Oe-circRNA group at 72 h (Figure 3B), suggesting that overexpression of hsa_circ_0068033 could induce apoptosis in MCF-7 and MDA-MB-231 cells. The immunoblotting test revealed that the expression of Bcl-2 protein was decreased and procaspase-3 splicing was activated (the expression of the spliceosome was increased accordingly) in the Oe-circRNA groups of the two cell lines (Figure 3C and 3D). These findings indicated that hsa_circ_0068033 may induce apoptosis in MCF-7 and MDA-MB-231 cells by activating the endogenous apoptotic pathway.

**Figure 3.**
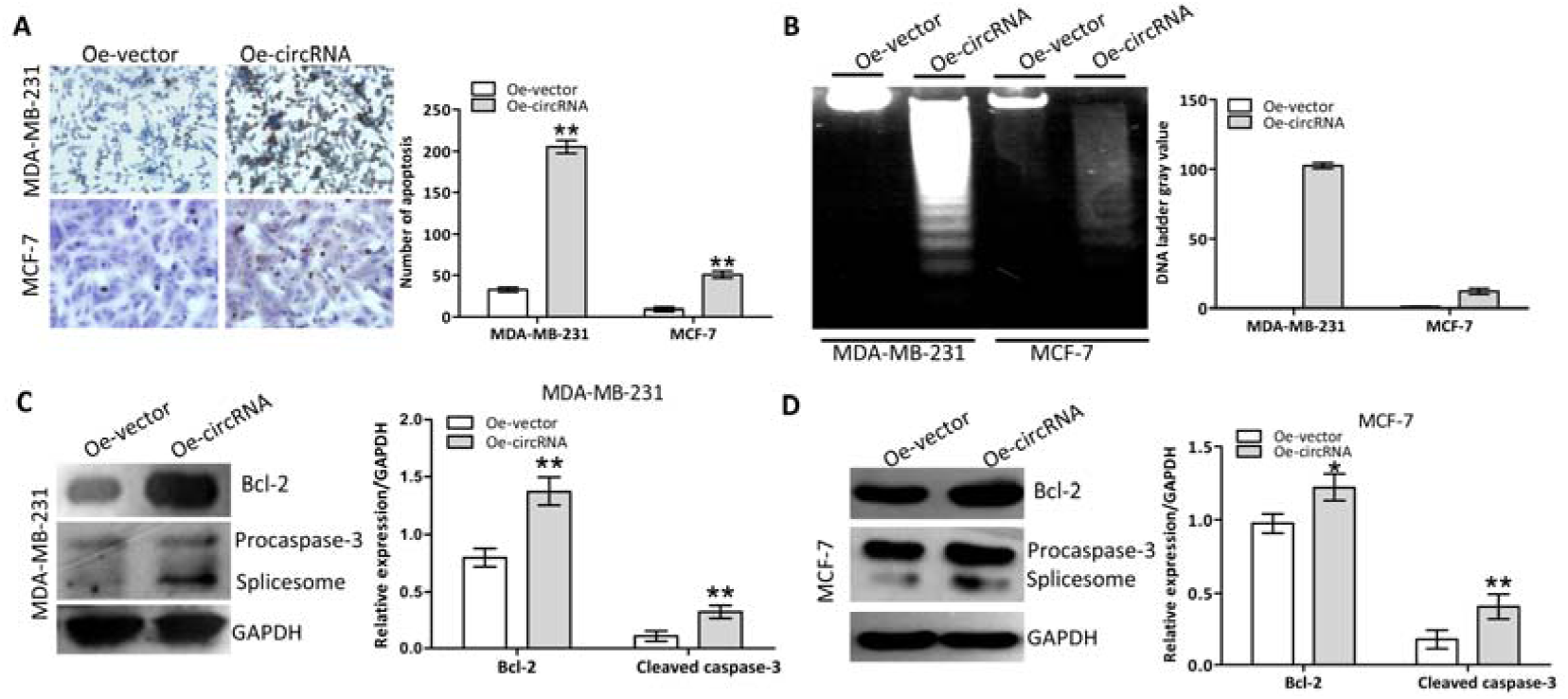
Overexpression of hsa_circ_0068033 induced apoptosis in MCF-7 and MDA-MB-231 cells. (A) The TUNEL assay showed increasing apoptosis in MCF-7 and MDA-MB-231 cells following the overexpression of hsa_circ_0068033. Apoptotic cells were counted 72 h following transfection. (B) DNA ladder assay. Images were captured 72 h following transfection with PLCDH-cirR0068033. (C, D) Immunoblotting showed activation of the intrinsic apoptotic proteins Bcl-2 and procaspase-3 in the two cell lines in response to overexpression of hsa_circ_0068033. \*\**P*<0.01.

### 3.4. Hsa_circ_0068033 regulated the progression of BC by acting as the sponge of miR-659

Hsa_circ_0068033 was mainly distributed in the cytoplasm of MCF-7 and MDA-MB-231 cells (Figure 4A). Thus, we speculated that hsa_circ_0068033 may play a regulatory role as a miRNA sponge. Based on the bioinformatics analysis, we predicted that 37 miRNA molecules may interact with hsa_circ_0068033, and miR-659 and miR-758 had the highest predicted scores (data not shown). The qRT-PCR analysis suggested that the expression of miR-659 in BC tissues was apparently higher than that reported in normal adjacent tissues (Figure 4B). This indicates a potential interaction between miR-659 and hsa_circ_0068033. Based on this evidence, we further designed a miR-mimic experiment. After transfection of the two cell lines with the miR-659 mimic, the expression of miR-659 was obviously increased (Figure 4C). Additionally, the dual luciferase reporter assay indicated that the luciferase activity in the cirRNA-wt group co-transfected with the miR-659 mimic in MDA-MB-231 cells was markedly decreased than that observed in the negative control group. In contrast, the luciferase activity in the cirRNA-mut group did not exhibit evident fluctuations (Figure 4D). The CCK-8, Transwell, and TUNEL experiments further revealed that the growth of MCF-7 and MDA-MB-231 cells was markedly faster (Figure 4E), their invasion and migration abilities were increased (Figure 4F and 4G), and the apoptosis-inducing effect was weaker (Figure 4H) in the Oe-circRNA+mimic group compared with the Oe-circRNA group.

**Figure 4.**
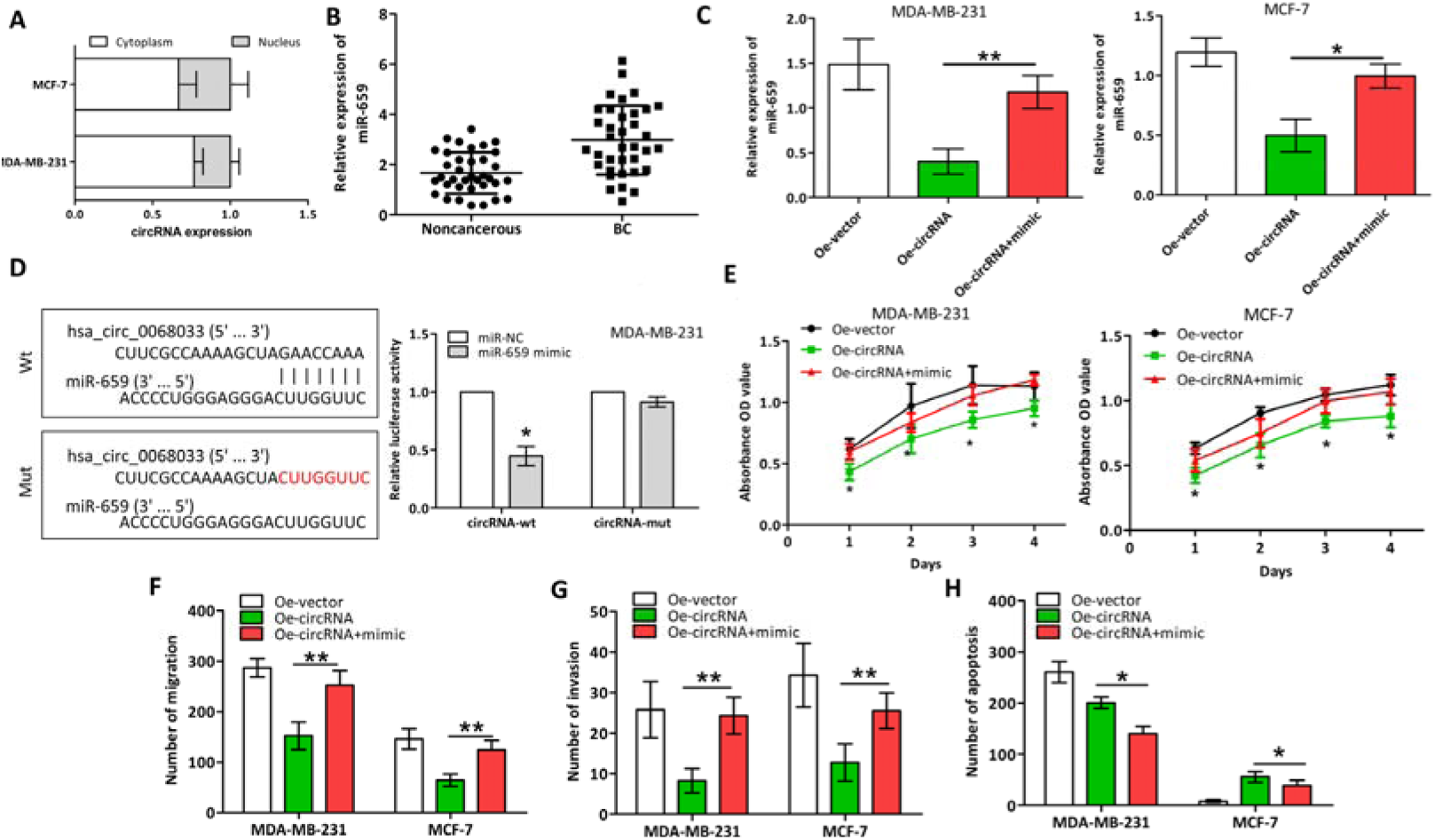
Hsa_circ_0068033 acts as a sponge of miR-659 to regulate the progression of BC. (A) Distribution of hsa_circ_0068033 in the nuclear and cytoplasmic fractions of MCF-7 and MDA-MB-231 cells analyzed through qRT-PCR. (B) The expression of hsa_circ_0068033 was upregulated in BC tissues compared with non-cancerous tissues. (C) MiR-659 levels in the two cell lines transfected with PLCDH-cirR0068033 and the miR-659 mimic. (D) Luciferase assay of MDA-MB-231 cells co-transfected with the miR-659 mimic and luciferase reporter containing hsa_circ_0068033-wild sequences (cirRNA-wt) or -mutant sequences (cirRNA-mut). (E) CCK-8 analysis. (F) Wound healing migration assay. (G) Transwell assay. (H) TUNEL apoptosis assay to verify the effects of hsa_circ_0068033 as a sponge of miR-659. \**P*<0.05, \*\**P*<0.01.

## 4. Discussion

CircRNA, a class of enclosed non-coding RNA molecules generated by RNA splicing, function as an miRNA sponge and regulate gene transcription and alternative splicing [18, 19, 8, 13, 4, 5]. Recent studies have confirmed that circRNA is involved in the regulation of gene expression by various means, thereby indirectly participating in the occurrence and development of various solid cancers [13, 11, 10, 6, 17, 9, 16, 14]. However, the biological functions of circRNA in BC are scarcely reported. In this study, we investigated the expression of hsa_circ_0068033 in BC tissues and cell lines and explored the influence of overexpressed hsa_circ_0068033 on the biological characteristics of MCF-7 and MDA-MB-231. In addition, we examined the potential molecular mechanism involved in this process through *in-vitro* and *in-vivo* experiments. We found that the expression of hsa_circ_0068033 was down-regulated in BC tissues and cell lines, while overexpression of hsa_circ_0068033 could inhibit the growth, proliferation, invasion, migration, and tumor formation of MCF-7 and MDA-MB-231 cells, and induce apoptosis.

Studies based on circRNA microarray analysis have shown that multiple circRNAs are abnormally expressed in BC [20, 21]. For example, a study performed by Tang et al. has shown that 1,155 abnormally expressed circRNAs (715 down-regulated and 440 up-regulated) are present in BC tissues [20]. A chip analysis conducted by Zhang et al. has identified 1,314 abnormally expressed circRNAs, the majority of which are derived from coding genes Rev3l, IGSF11, MAML2, and LPP [21]. In this study, qRT-PCR and ISH confirmed that the expression of hsa_circ_0068033 was decreased in BC tissues and four BC cell lines, and this expression was associated with the clinical staging of patients. This suggests that hsa_circ_0068033 may participate in the progression of BC. Moreover, ROC analysis suggested a high diagnostic value of hsa_circ_0068033 (AUC=0.8480), implying that it may become a new marker for the diagnosis of BC.

CircRNAs probably have different biological functions in BC. Tang et al. discovered that highly expressed hsa_circ_0001982 can inhibit the growth, proliferation, invasion, and migration of tumor cells, and induce their apoptosis [20]. Another study reported that the expression of circVRK1 is evidently down-regulated, and circVRK1 functions as a cancer suppressor gene [22]. Similarly, the present study found that exogenous hsa_circ_0068033 could significantly inhibit the growth, clone formation, invasion, and migration of MCF-7 and MDA-MB-231 cells, and induce apoptosis in BC cells by activating endogenous mitochondrial pathways. This indicates that hsa_circ_0068033 acts as a cancer suppressor gene in BC, which is consistent with our clinical results. This study further verified our hypothesis through *in-vivo* experiments. Overexpression of hsa_circ_0068033 significantly inhibited tumor formation in MDA-MB-231 cells. Intriguingly, there was no tumor formation in the Oe-circRNA and Oe-vector groups after injection of MCF-7 cells. This is possibly associated with the characteristics of MCF-7 cells we cultured.

Studies have shown that certain types of circRNAs, such as hsa_circ_0001982 [20], circGFRA1 [23], circ-ABCB10 [24], circ-Foxo3 [25], and circRNA_1093[21], may function as a miRNA sponge in BC cells. Our qRT-PCR analysis found that hsa_circ_0068033 was mainly distributed in the cytoplasm of MDA-MB-231 and MCF-7 cells, suggesting that it may play the role of a miRNA sponge. With the assistance of the bioinformatics analysis, we predicted that 37 miRNA molecules may interact with hsa_circ_0068033. Notably, miR-659 and miR-758 had the highest prediction scores. Through the qRT-PCR analysis, we verified that miR-659 exhibits a higher expression kurtosis than miR-758 in BC tissues. Our subsequent investigation further revealed that the miR-659 mimic was able to reverse the tumor-suppressing and apoptosis-inducing effect of hsa_circ_0068033 in MCF-7 and MDA-MB-231 cells. Moreover, we further confirmed the interaction between hsa_circ_0068033 and miR-659.

Nevertheless, this study has some shortcomings. Firstly, only 36 clinical cases with BC were included; hence, the sample size was small. Secondly, we only performed TUNEL and DNA ladder experiments, and merely examined a few intrinsic apoptotic proteins in the study of the molecular mechanism of apoptosis. Thirdly, we failed to prove that hsa_circ_0068033 can serve as a miR-659 sponge to trigger interaction at the animal level. Furthermore, whether hsa_circ_0068033 interacts with other miRNAs warrants further investigation.

In conclusion, this study found that the expression of hsa_circ_0068033 is decreased in BC and may be associated with the progression of disease. Moreover, it was shown that hsa_circ_0068033 may be highly valuable in the diagnosis of BC. Moreover, the overexpression of hsa_circ_0068033 may inhibit the growth, proliferation, invasion, migration, and tumor formation of BC cells, and induce apoptosis by activating endogenous apoptotic pathways. Lastly, hsa_circ_0068033 may participate in the progression of BC as a miR-659 sponge.

## Data Availability Statement

The original data used to support the findings of this study are available from the corresponding author upon request.

## Author contributions

Conception and design: DC Liu; Administrative support: DC Liu Chang; Provision of study materials or patients: PF Yuan; Collection and assembly of data: PF Yuan and LL Lei; Data analysis and interpretation: PF Yuan and LL Lei; Manuscript writing: All authors; Final approval of manuscript: All authors.

## Conflicts of Interest

The authors declare no conflicts of interest.

## Acknowledgments

This study was supported by the Henan Provincial Programs for Science and Technology Development (No.201504021).

## References

[1] R.L. Siegel, K.D. Miller, A. Jemal, Cancer statistics, 2016, CA: a cancer journal for clinicians 66 (2016) 7–30. 10.3322/caac.21332.

[2] R.L. Siegel, K.D. Miller, A. Jemal, Cancer statistics, 2019, CA: a cancer journal for clinicians 69 (2019) 7–34. 10.3322/caac.21551.

[3] L.J. Al-Mansouri, M.S. Alokail, Molecular basis of breast cancer, Saudi medical journal 27 (2006) 9–16.

[4] L.L. Chen, The biogenesis and emerging roles of circular RNAs, Nature reviews. Molecular cell biology 17 (2016) 205–211. 10.1038/nrm.2015.32.

[5] L.L. Chen, L. Yang, Regulation of circRNA biogenesis, RNA biology 12 (2015) 381–388. 10.1080/15476286.2015.1020271.

[6] S. Qu, Z. Liu, X. Yang, J. Zhou, H. Yu, R. Zhang, H. Li, The emerging functions and roles of circular RNAs in cancer, Cancer letters 414 (2018) 301–309. 10.1016/j.canlet.2017.11.022.

[7] Y. Dong, D. He, Z. Peng, W. Peng, W. Shi, J. Wang, B. Li, C. Zhang, C. Duan, Circular RNAs in cancer: an emerging key player, Journal of hematology & oncology 10 (2017) 2. 10.1186/s13045-016-0370-2.

[8] D.H. Bach, S.K. Lee, A.K. Sood, Circular RNAs in Cancer, Molecular therapy. Nucleic acids 16 (2019) 118–129. 10.1016/j.omtn.2019.02.005.

[9] M. Song, L. Xia, M. Sun, C. Yang, F. Wang, Circular RNA in Liver: Health and Diseases, Advances in experimental medicine and biology 1087 (2018) 245–257. 10.1007/978-981-13-1426-1_20.

[10] W.L. Ng, T.B. Mohd Mohidin, K. Shukla, Functional role of circular RNAs in cancer development and progression, RNA biology 15 (2018) 995–1005. 10.1080/15476286.2018.1486659.

[11] Y. Chen, S. Wei, X. Wang, X. Zhu, S. Han, Progress in research on the role of circular RNAs in lung cancer, World journal of surgical oncology 16 (2018) 215. 10.1186/s12957-018-1515-2.

[12] X. Di, X. Jin, R. Li, M. Zhao, K. Wang, CircRNAs and lung cancer: Biomarkers and master regulators, Life sciences 220 (2019) 177–185. 10.1016/j.lfs.2019.01.055.

[13] S. Chaichian, R. Shafabakhsh, S.M. Mirhashemi, B. Moazzami, Z. Asemi, Circular RNAs: A novel biomarker for cervical cancer, Journal of cellular physiology (2019). 10.1002/jcp.29009.

[14] K.W. Wang, M. Dong, Role of circular RNAs in gastric cancer: Recent advances and prospects, World journal of gastrointestinal oncology 11 (2019) 459–469. 10.4251/wjgo.v11.i6.459.

[15] J. Li, J. Yang, P. Zhou, Y. Le, C. Zhou, S. Wang, D. Xu, H.K. Lin, Z. Gong, Circular RNAs in cancer: novel insights into origins, properties, functions and implications, American journal of cancer research 5 (2015) 472–480.

[16] M.I. Taborda, S. Ramirez, G. Bernal, Circular RNAs in colorectal cancer: Possible roles in regulation of cancer cells, World journal of gastrointestinal oncology 9 (2017) 62–69. 10.4251/wjgo.v9.i2.62.

[17] H. Shuwen, Z. Qing, Z. Yan, Y. Xi, Competitive endogenous RNA in colorectal cancer: A systematic review, Gene 645 (2018) 157–162. 10.1016/j.gene.2017.12.036.

[18] H.M. Li, X.L. Ma, H.G. Li, Intriguing circles: Conflicts and controversies in circular RNA research, Wiley interdisciplinary reviews. RNA (2019) e1538. 10.1002/wrna.1538.

[19] A.C. Panda, Circular RNAs Act as miRNA Sponges, Advances in experimental medicine and biology 1087 (2018) 67–79. 10.1007/978-981-13-1426-1_6.

[20] Y.Y. Tang, P. Zhao, T.N. Zou, J.J. Duan, R. Zhi, S.Y. Yang, D.C. Yang, X.L. Wang, Circular RNA hsa_circ_0001982 Promotes Breast Cancer Cell Carcinogenesis Through Decreasing miR-143, DNA and cell biology 36 (2017) 901–908. 10.1089/dna.2017.3862.

[21] C. Zhang, H. Wu, Y. Wang, Y. Zhao, X. Fang, C. Chen, H. Chen, Expression Patterns of Circular RNAs from Primary Kinase Transcripts in the Mammary Glands of Lactating Rats, Journal of breast cancer 18 (2015) 235–241. 10.4048/jbc.2015.18.3.235.

[22] N. Yan, H. Xu, J. Zhang, L. Xu, Y. Zhang, L. Zhang, Y. Xu, F. Zhang, Circular RNA profile indicates circular RNA VRK1 is negatively related with breast cancer stem cells, Oncotarget 8 (2017) 95704–95718. 10.18632/oncotarget.21183.

[23] R. He, P. Liu, X. Xie, Y. Zhou, Q. Liao, W. Xiong, X. Li, G. Li, Z. Zeng, H. Tang, circGFRA1 and GFRA1 act as ceRNAs in triple negative breast cancer by regulating miR-34a, Journal of experimental & clinical cancer research : CR 36 (2017) 145. 10.1186/s13046-017-0614-1.

[24] H.F. Liang, X.Z. Zhang, B.G. Liu, G.T. Jia, W.L. Li, Circular RNA circ-ABCB10 promotes breast cancer proliferation and progression through sponging miR-1271, American journal of cancer research 7 (2017) 1566–1576.

[25] W. Yang, W.W. Du, X. Li, A.J. Yee, B.B. Yang, Foxo3 activity promoted by non-coding effects of circular RNA and Foxo3 pseudogene in the inhibition of tumor growth and angiogenesis, Oncogene 35 (2016) 3919–3931. 10.1038/onc.2015.460.

